# Avian promiscuity in the last European primeval forest

**DOI:** 10.1101/2025.05.29.656776

**Authors:** Joanna Sudyka, Irene Di Lecce, Grzegorz Hebda, Patryk Rowiński, Robert Rutkowski, Anna Szczuka, Charles Perrier, Marta Cholewa, Marta Maziarz, Tomasz Wesołowski, Marta Szulkin

## Abstract

Primeval forests provide a reference baseline to understand the origins and evolution of mating systems because they offer a unique, undisturbed environment where ecological interactions and natural selection play out in their original context. We established rates of extra-pair paternity (EPP) for two bird species referential to evolutionary ecology: blue tits (*Cyanistes caeruleus*) and great tits (*Parus major*), breeding in natural cavities in Białowieża National Park, the sole remnant of European lowland primeval forests. Genotype-by-sequencing of 889 blue tits and 1,256 great tits revealed that 48% of blue tit broods and 39% great tit broods were of mixed paternity, with 15% blue tit and 14% great tit nestlings not sired by their social father. These referential levels align with median values of EPP reported in nestbox studies in secondary and managed forests, suggesting that certain reproductive strategies are advantageous across diverse environments. Observed promiscuity levels did not affect reproductive success or parental investment, indicating no or limited adaptive value for promiscuity in the context of a primeval forest ecosystem. Our study underscores how primeval habitats provide unparalleled insights into natural selection and mating strategies in wild populations, highlighting the stability and resilience of these strategies across different environmental contexts.

## Introduction

Primeval forests, characterized by long-term ecological continuity free from major direct human disturbance, are a rarity across the globe. In Europe, these forests are even scarcer and contribute only 0.4% to the global area of primary forests^1^. Białowieża Forest is the sole iconic remnant of a mixed forest that once stretched across Central Europe. The area’s unique history allowed its protection through the ages, long before large-scale anthropopressure^2^. The role of such ecosystems is irreplaceable, showing resilience to non-native species invasion^3^ and contributing to global climate and biodiversity targets^3^. Ecological networks inherent to Białowieża Forest are our best reference baseline to a specific biome of origin for many native European plant and animal species. These species further exercise their plasticity and adaptability in anthropogenically altered habitats, ranging from secondary forests to urbanized areas. While the fundamental importance of intact landscapes is much discussed in biodiversity and community ecology^4^, knowledge on mating systems and sexual selection, the driving forces of adaptation and evolution, remains understudied in these ecosystems. Much of what we know about mating strategies, which are deeply intertwined with ecological interactions such as predator-prey dynamics and resource/breeding site competition, comes from studies carried out in more accessible or impacted ecosystems, such as managed forests and fragmented habitats, or in controlled environments such as enclosures or laboratories. However, modified habitats with extensive fragmentation and various degrees of anthropogenic disturbance can alter mating behavior, a phenomenon well documented in avian species^5,6^. It thus remains unresolved if promiscuity rates among socially monogamous birds reported to date, where on average 11% of offspring have an extra-pair father^7^, represent the natural benchmark for adaptation and evolution of mating strategies.

Promiscuous mating can benefit males, who increase their contribution to the gene pool, and females, who secure optimal genes for their offspring^8^. These functional insights are mainly derived from avian populations breeding in secondary and managed forests, thereby experiencing altered breeding ecology compared to natural populations that evolved and optimized their reproduction in natural cavities of primeval forests. Nestbox provisioning in itself is another human impact on ecosystems, as it may increase breeding densities, alter predation pressures, and modify reproductive success^9–13^. Promiscuity may increase with the density of breeding pairs^14^, and the opportunity to engage in extra-pair copulations is greater in anthropogenically altered habitats^5^, leading us to expect lower promiscuity levels in unaltered primeval habitats.

In this study, we examined extra-pair paternity (EPP) of two model species for avian promiscuity^15^ in natural cavities of Białowieża primeval forest (Fig. S1). Specifically, we genotyped 889 blue tits (*Cyanistes caeruleus*) at 14,019 SNP markers and 1,256 great tits (*Parus major*) at 13,754 SNP markers with the aim of establishing reference baseline levels for promiscuity and assessing the applicability of existing nestbox-based ecological models that quantify promiscuity levels in natural populations. We further empirically tested how phenological and environmental parameters impacted promiscuity levels derived from natural cavities to uncover the ecological and evolutionary drivers of mating strategies under primeval conditions. Finally, we assessed whether promiscuity covaried with reproductive success and parental investment in an effort to establish its broad adaptive value and understand natural selection in this complex ecosystem.

## Materials and methods

### Study site, fieldwork and sampling

Białowieża Forest, located in the heart of the European plain along the Polish-Belarusian border (Białowieża village: 52°41′ N, 23°52′ E), is an unparalleled natural treasure. It is the last remaining fragment of the vast lowland forests that once covered much of temperate Europe, distinguished by its remarkable size and exceptional state of preservation^16–18^. Most of its 613 km^2^ Polish section (about 45% of the total area) is managed, yet a 48 km^2^ (4,800 ha) block of primeval old-growth forest within Białowieża National Park (BNP) remains strictly protected. This pristine forest is a multistorey, mixed-species ecosystem with uneven-aged stands featuring numerous veteran trees, including the Norway spruce (*Picea abies*), up to 52 meters high^19^. Unmanaged deadwood accounts for 20–25% of the total wood volume, fostering exceptional biodiversity^19^. Białowieża Forest is a singular relic of Europe’s reference biome, embodying a level of ecological integrity that is unmatched anywhere else on the continent. Fieldwork was conducted in four large plots (33–54 ha each) within BNP, totaling 185 ha and spaced 1–2 km apart (Fig. S1). Three plots (C, M, W) were in oak-lime-hornbeam stands dominated by hornbeam (*Carpinus betulus*), small-leaved lime (*Tilia cordata*), pedunculate oak (*Quercus robur*), spruce (*Picea abies*), and continental maple (*Acer platanoides*)^20,21^. The average breeding pair density per 10 ha in the oak–hornbeam–lime forest patches in BNP was 4.0 pairs for blue tits and 4.9 pairs for great tits^18^. The fourth plot (K), with similar densities, was in a swampy riverine forest of alder (*Alnus glutinosa*), ash (*Fraxinus excelsior*), and spruce, and birds breeding in plot K had access to nearby drier hornbeam-covered areas^16^. Natural cavities were abundant in all plots, and tits exclusively used these for breeding^17^. Intensive nest searches were conducted to locate all natural cavities in two of the oak-lime-hornbeam areas: plot C (48 ha) and plot M (54 ha)^20,21^. Additional nest searches were conducted less intensively during general mapping censuses in plots W (50 ha) and K (33 ha)^19^.

Blue tit DNA samples were collected between 2005 and 2008^19,21^, and great tit DNA samples were collected between 2007 and 2011^20^ (Table S1). Working with passerines in natural cavities is more complex and laborious compared to nestboxes^11^. Even before breeding began, birds were tracked and caught from winter (November, December) till early spring (beginning of April). Birds were uniquely color-ringed, aged as 1^st^ year breeders or older based on wing molt patterns, and their movements were recorded on field maps to estimate the number and distribution of breeding pairs, simplifying mapping and nest discovery. Natural cavities used for breeding were found by following birds mostly during nest-building and were regularly monitored to establish brood productivity and nest fate (failed or successful), with visit frequency adjusted based on the nesting stage^20,21^. Lower cavities (up to 6 m) were inspected from the ground or with a ladder, while higher ones by climbing with special tree-climbing spikes. This allowed to acquire data on lay date, clutch size, hatch date, number of hatchlings, fledging date, number of successfully fledged nestlings, and fledging age of nestlings (i.e. duration of the nestling period, from hatching till fledging). An array of environmental variables that precisely described each cavity was also collected, such as cavity height, entrance and nest chamber dimensions, cavity exposure (see Supplementary Material: “*Phenological and environmental variables”* and Figs. S2 and S3 for a complete list of variables and their correlations). The effect of these phenological and environmental variables was further tested against promiscuity rates. The number of fledged nestlings and fledging age were treated as best proxies for reproductive success and parental investment obtained in our study system. Both variables were tested for a covariation with brood promiscuity levels. Twelve days after hatching, two tail feathers were collected and immediately stored in 96% ethanol from (i) nestlings safely removed from nests using a custom-made loop tool and (ii) from parents captured using mist nets or traps near nest entrances (if they had not been captured before, i.e., they were unringed). All nestlings were ringed with a uniquely numbered metal leg band.

### Genetic analysis and parentage assignment

Blue tit feather DNA was extracted with QIAamp DNA Investigator Kit (adults) and QuickGene DNA tissue kit (nestlings; Qiagen, Hilden, Germany). Great tit samples were extracted with Quick-DNA™ Miniprep Plus Kit (Zymo Research) according to the manufacturer’s protocol. A section of the feather interior (shaft’s superior umbilicus), where the blood clot is located, was cut into small pieces (2 mm each) for overnight lysis at 38°C. DNA sequencing was performed by Diversity Arrays Technology Pty, Ltd (Canberra, AU) using DArTseqLD, a high-throughput genotyping by sequencing method that employs genomic complexity reduction using restriction enzyme pairs^22^. In blue tits, we obtained 908 samples genotyped at 14,019 SNP markers and in great tits, 1,286 samples genotyped at 13,754 SNP markers. Details on the DArT sequencing technology can be found in^5,10^. All subsequent analyses were run separately for each species in R (v. 4.3.2)^23^. Using dartR (v. 2.9.7)^24^, we filtered out individuals with call rates below 20% and loci with call rates below 40% in blue tits, and both individuals and loci with call rates below 40% in great tits. These thresholds were selected in order to retain as much information as possible, given the low call rate in adults (Fig. S4). After filtering, 12,248 SNP markers and 884 individuals were kept in blue tits (889 individuals when including five recruits, whose genotypes were used twice: first to determine their EPO status within their brood of rearing, and later as parental genotypes in subsequent breeding seasons), and 12,191 SNP markers and 1,256 individuals were kept in great tits. We estimated genetic relationships among individuals using the function *snpgdsGRM* with the method GCTA^25^ implemented in SNPRelate (version 1.36.0)^26^. *snpgdsGRM* was run with a minor allele frequency threshold of 0.005 and a missing rate of 0.05 in both species. A total of 2,136 loci were used in blue tits and a total of 3,711 loci were used in great tits. We confirmed that the effect of using these low filtering thresholds on genetic relatedness was negligible, by testing it on a subset of individuals with high call rate: relatedness was calculated using 40% of the SNP data and compared to relatedness calculated with 100% of the SNP data (Fig. S5). We compared the resulting genomewide relatedness matrix (GRM) with a social pedigree of all individuals ringed in the field created using *ggroups* (v. 2.1.0)^27^. Following Perrier et al.^28^ and Di Lecce et al.^5^, discrepancies between the GRM and the social pedigree were identified separately among (i) mothers with their offspring, (ii) social fathers with their offspring and (iii) offspring within their own nest, in nests where the social father was not sampled. Father– offspring pairs (social relatedness = 0.5) showing GRM relatedness estimates below 0.3 (based on relationships between mothers and their offspring; Fig. S6a) were classified as instances of EPP. In nests where the social father or both social parents were not sampled (e.g. when catching attempts failed or when a parent abandoned the nest – 37% of nests in blue tits and 7% in great tits), we did not assign within-pair or EPP status to individual nestlings but only recorded whether the brood was of mixed-paternity, that is, whether EPP occurred (these nests were included in all analyses on EPP occurrence, but not on proportion of extra-pair offspring - EPO). Therefore, if there were pairs of siblings within a given nest (social relatedness = 0.5) with GRM estimates between 0.15 and 0.35, the nest was classified as containing half-siblings; if all pairs of siblings within a given nest had GRM estimates above 0.35, the nest was classified as containing full-siblings. Nestlings with GRM relatedness estimates below 0.15 with both the social parents and the social siblings (social relatedness = 0.5) were classified as instances of brood parasitism (Fig. S6). The SNP parentage assignment was additionally validated with a microsatellite parentage analysis in blue tits, and the results were highly consistent (Supplementary Material: *Microsatellite parentage analysis and comparison with SNP data*). Overall, the occurrence of mixed paternity in a brood was highly consistent between the methods (McNemar’s test comparing EPP detected by microsatellites vs. SNPs: χ^2^ = 0, df = 1, p = 1), and the relationship between the number of offspring assigned as extra-pair per brood was strong between the methods (Spearman’s rho = 0.816, p < 0.0001).

### Statistical analyses

To examine the role of environmental and phenological breeding parameters on promiscuity, we fitted EPP occurrence (0 if none of the nestlings was extra-pair or 1 if at least one nestling was extra-pair in a nest) and EPO proportion (the number of EPO over the number of genotyped nestlings per brood, the ratio coded as a continuous value between 0 and 1) as dependent variables in two separate generalized linear mixed models (GLMMs) for each species separately in R (v. 4.2.2)^29^. As there was mortality recorded between hatching and time of sampling, and parents removed dead nestlings early on during development, we included all broods in which we successfully sampled and genotyped at least 50% of the original number of hatched nestlings, in order to gain a representative sample size of the original brood in all analyses involving EPP occurrence and EPO proportion (8 nests in blue tits and 6 in great tits were removed). Each model was run with a binomial distribution and logit link function, and the models on EPO proportion were additionally weighed for the number of genotyped nestlings per brood. To avoid multicollinearity, reduce model overfitting, and improve the reliability of model selection, we included only one variable from six sets of highly correlated phenological and environmental predictors. Each set informed of different environmental gradients (Supplementary Material: *Phenological and environmental variables*; Figs. S2 and S3). Within each set, we retained the variable recorded in the highest number of nests. Specifically, we selected: lay date, initial brood size, height, entrance area, depth and nest bottom area. These six retained variables were then used as continuous predictors in model selection procedures, to which we also included nest exposure (coolest, intermediate, warmest) as a fixed predictor. The random effects structure included year and study plot (C, M, which had the most nests located, and “Other” - encompassing study plots W and K with fewer nests present, to balance out the sample sizes). We then performed AIC-based model selection with *dredge()* function of the MuMIn package^30^, as we did not have specific assumptions to test which of the variables would contribute most to variance in promiscuity.

To test the relationship between promiscuity and available proxies of reproductive success and parental investment, we fitted separate linear mixed models (LMMs) per species, with responses defined as the number of nestlings that successfully fledged (a universal measure of avian reproductive success^31^) and nestling fledging age (an indicator of parental investment: indeed, longer nesting periods may enhance juvenile survival through increased parental care^32^, while also increase predation risk^33^, especially in a primeval environment where there is an abundance of predators^17^). In models on the number of fledged nestlings in great tits, more nests failed than in blue tits - thus, great tit fledgling number was more often 0. To model great tit reproductive success, we therefore used a GLMM with poisson distribution as it provided a better fit, and lower AIC compared to fitting a separate zero-inflation process. We again fitted the same set of seven predictors, i.e., lay date, initial brood size, height, entrance area, depth, nest bottom area and exposure, alongside plot and year as random effects. Here, we fitted separate models, testing two promiscuity-related predictors to assess the qualitative (EPP occurrence - fixed) and quantitative (EPO proportion - continuous) contributors to reproductive success. We used the *fixed* condition in *dredge()* function of the MuMIn package^30^ to ensure these were always retained as variables of interest.

To identify the most parsimonious model, we performed model selection using AICc-based model averaging (*MuMIn::dredge*). Predictors appearing in models with ΔAICc <1.5 (rather than 2 as usually applied^34^ to reduce model uncertainty and to include fewer meaningful predictors from more robust models) were retained and refitted in a single linear mixed-effects model to obtain parameter estimates, random effect variance, multicollinearity diagnostics (VIF), and marginal/conditional R^2^. Due to problems with a singular fit at some models’ selection stage, we removed random effects for which the variance was zero or close to zero, before running *dredge()*. We calculated *R*^2^ as defined in^35^ for each model with the MuMIn package^30^. We tested all final models for multicollinearity (VIF scores never exceeded 2). We checked each model fit (KS test), dispersion and identified potential outliers by simulating residuals and plotting residual diagnostics using the DHARMa package^36^. Outliers were identified and tested by re-running the models on adjusted datasets. In all cases, removing outliers did not alter the factor estimates either qualitatively or quantitatively, so all outliers were retained. The only exception was the model examining the proportion of EPO in great tits, where outliers did affect model estimates. However, these values fell within the expected biological range and were not highly influential. Therefore, we chose to retain them in order to preserve ecological validity.

## Results and discussion

This first report on EPP in two model avian species breeding in natural cavities of a primeval forest - a reference biome for studying long-term evolutionary processes unaffected by recent anthropogenic disturbances – establishes baseline promiscuity levels that are instrumental for the study of mating strategies. We report that EPP in blue tits was recorded in 48% of all broods (46 out of 96 broods were of mixed-paternity), with 15% of offspring being extra-pair (56 nestlings out of 362). In great tits, EPP was recorded in 39% of broods (47 out of 122 broods were of mixed paternity), with 14% of EPO (111 nestlings out of 821) (Fig.1a). These levels are in line with values reported in more commonly studied environments – nestbox populations of blue tits and great tits breeding in secondary forests (Fig. 1b).

**Figure 1.**
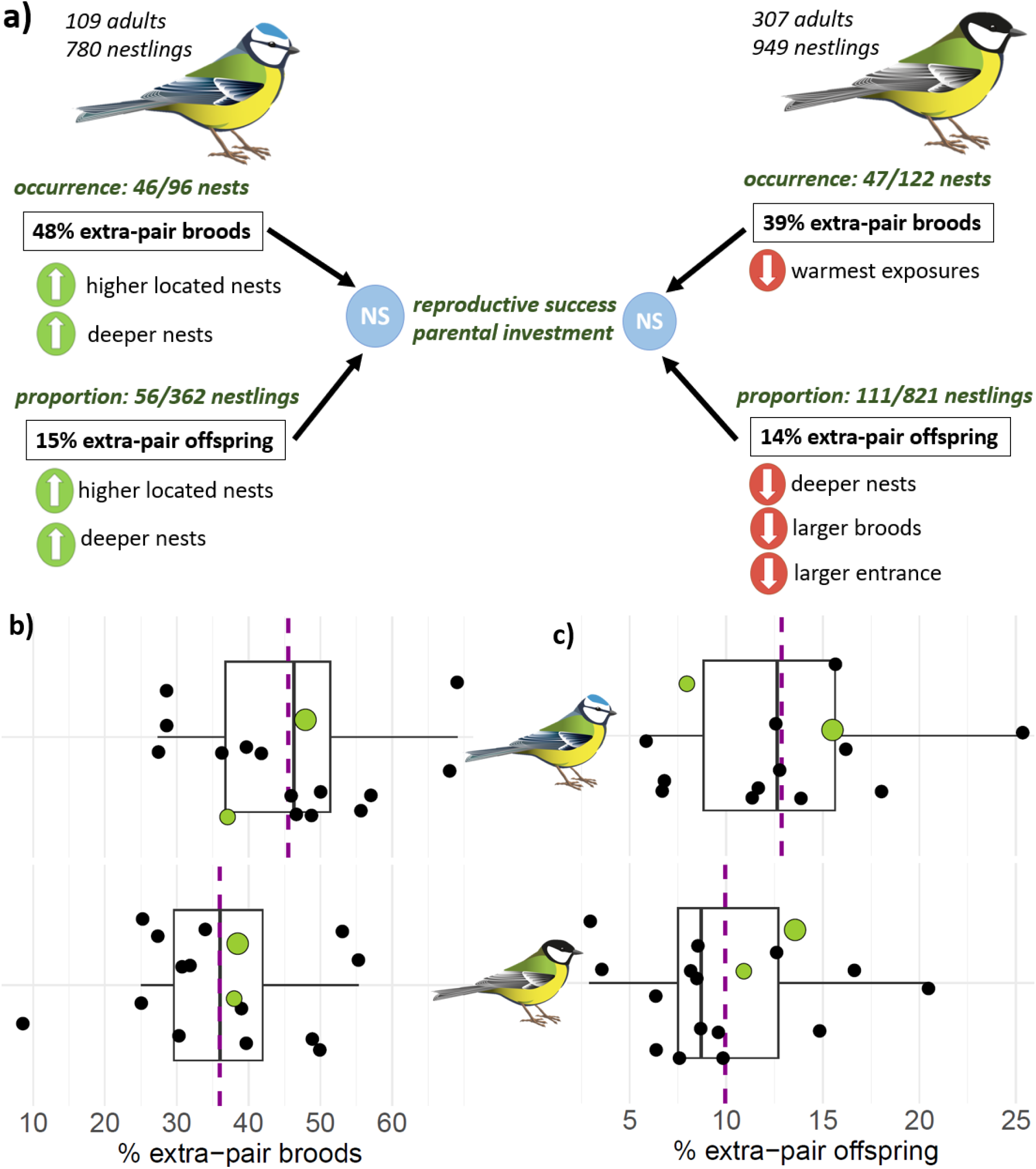
**a)** Primeval forest EPP occurrence and proportion of EPO in the brood in blue tits (left) and great tits (right), alongside their environmental and phenological contributors. There was no relationship between EPP occurrence or proportion of EPO in the brood and reproductive success (number of fledged offspring) or parental investment (age of nestlings at fledging) in Białowieża primeval forest. Green upside arrows denote positive correlations, red downward arrows - negative correlations, and blue NS signs - no correlation (Tables S2-S5). Percentages of **b)** extra-pair broods, with recorded EPP occurrence and **c)** EPO for blue tits (upper panels) and great tits (lower panels) reported across studies [black dots; Brouwer & Griffith (2019)^15^]. Values from primeval forest natural cavities are reported as large green dots. Values from the only other known natural cavity study^11^ are reported as small green dots. The magenta dashed line represents average values, while the box represents the median with IQR and whiskers representing 1.5 IQR.

To contextualize the values observed in Białowieża, we derived population-specific values of EPP in blue tits and great tits from a recent review by Brouwer & Griffith^15^, in which both species were by far the most studied avian model system in terms of promiscuity (9.5 k blue tit and 14 k great tit individuals assessed). We also added promiscuity values from a recent nestbox-natural cavity comparison to reference the only other study carried out in natural cavities so far^11^ (Fig. 1b). Since promiscuity values for both species reported from Białowieża fall within the middle of distributions for extra-pair broods and EPO ranges of the otherwise unexplained variation in promiscuity^15^, they likely represent the natural reference optimum advantageous across a range of environmental conditions, pointing to the evolutionary stability of mating systems. This finding validates and supports foundational evolutionary biology principles, suggesting that certain reproductive strategies are advantageous across diverse environments with a degree of behavioral plasticity. It affirms the universality of specific selection pressures (e.g., sexual selection, mate choice, or territorial behavior), extending the generalizability of ecological models. Future studies can thus use this primeval forest baseline to estimate the magnitude of shifts in promiscuity relative to natural levels.

Remarkably, observed variation in promiscuity levels in a primeval forest did not affect reproductive success or parental investment (Fig. 1a, Table S4 and S5). This suggests no or limited adaptive value and neutral selection for promiscuity in the context of a primeval forest ecosystem. In Białowieża, this might stem from high pressures by various predators (in particular forest dormouse, great spotted woodpecker, yellow-necked mouse, pine marten and weasel^21^), which are largely diminished in secondary forests or in nestbox populations^17^. Promiscuity may be here at equilibrium since there is no fitness advantage to altering its levels when predation is the major cause for reproductive failure - it accounts for 80% of nest failures in blue tits^21^ and 69% in great tits^20^. Despite demonstrated genetic benefits of promiscuity^8^, promiscuous copulations, competition for partners and mate guarding might attract predators to the nest and increase energy expenditure or time spent away from more defensive or survival-focused activities^15^. If the environment is heavily influenced by predation, fitness could rely more heavily on survival than on reproduction alone. Since in Białowieża primeval forest, promiscuity was not affecting reproductive success or parental investment directly (Fig. 1a, Table S4 and S5), it likely represented the optimal level, where individuals maximize their reproductive success while minimizing the time they spend exposed to predators. The importance of minimizing predation was also confirmed by lower numbers of fledged blue tits from cavities that had larger nest entrance areas, which were easier to depredate (Table S5A).

In general, we show lower levels of promiscuity in higher-quality nests: safer from predators (lower above the ground and deeper in a tree cavity), with greater reproductive investment (large broods) and better thermal properties (warmer exposures) (Fig. 1a). Specifically, in blue tits, promiscuity increased in terms of occurrence and offspring proportion in higher-located nests (Table S2 and S3). Higher nests may be harder for the social father to guard, but more importantly, increasing height has been previously demonstrated to be the largest negative contributor to nesting success in blue tits, as high nests were exposed to higher predation risk^21^. In great tits, we observed less promiscuity in deeper nests (Table S3), which were previously shown to be more successful and harder to predate^20^ and in warmer exposures (Table S2), with favorable microclimate that allows to reduce costs of incubation and nestling thermoregulation^37^. Moreover, we observed less promiscuity in nests with larger entrances (Table S3), and entrance area was also shown to contribute positively to breeding success in Białowieża great tits^20^. This reduction of promiscuity in good-quality nests can be readily explained by prioritization of the current reproductive success over promiscuity-related risks and has been reported previously^38–40^. Interestingly, in deeper blue tit nests, which also tend to be safer^21^, there was more promiscuity (Table S2 and S3). The divergent results with respect to nest depth between the species are difficult to explain, but nest depth is a much stronger factor in nesting success for great tits, whose nests are generally deeper than those of blue tits^20,21^. Different predator pressures may therefore influence how depth contributes to nest quality perception in each species.

Ultimately, showing that mating strategies in primeval forests align with those observed elsewhere highlights the stability and resilience of these strategies across different environmental contexts, impacting both the theory and application of evolutionary biology and ecology.

## Ethics approval

All methods and sampling detailed in this manuscript were performed under the approval of the Local Ethical Committee under Warsaw University of Life Sciences, SGGW, no. 50/2002, and the Directorate of Białowieża National Park dated 19/04/2005.

## Acknowledgments

We thank members of the Białowieża fieldwork team and numerous field assistants for their contribution to data collection. We also thank Hannah Dugdale for her support in data analysis.

## Author contributions

JS, GH, PR, TW and MS conceived and designed the study. GH, PR, MC, MM and TW performed the fieldwork. JS, RR and AS performed lab analyses. IDL and CP performed parentage assignment, and JS performed the statistical data analysis. JS wrote the manuscript with input from all authors. All authors approved the final version of the manuscript.

## Data and materials availability

The R code used in this study and raw data can be found at https://zenodo.org/records/15487049^41^.

