## Supplementary Material for "Avian promiscuity in the last European primeval forest"

### Authors' list and affiliations

Joanna Sudyka[1,2*], Irene Di Lecce[3], Grzegorz Hebda[4], Patryk Rowiński[5], Robert Rutkowski[6], Anna Szczuka[7], Charles Perrier[8], Marta Cholewa[9], Marta Maziarz[6], Tomasz Wesołowski[9] and Marta Szulkin[3*]

[1] Institute of Environmental Sciences, Jagiellonian University, Poland

[2] Groningen Institute for Evolutionary Life Sciences (GELIFES), Netherlands

[3] Institute of Evolutionary Biology, Faculty of Biology, University of Warsaw, Poland

[4] Institute of Biology, Opole University, Opole, Poland

[5] Department of Forest Zoology and Wildlife Management, Institute of Forest Science, Warsaw University of Life Sciences, SGGW, Poland;

[6] Museum and Institute of Zoology, Polish Academy of Sciences, Poland;

[7] Laboratory of Ethology, Nencki Institute of Experimental Biology, Polish Academy of Sciences, Poland;

[8] CBGP, INRAe, CIRAD, IRD, Montpellier SupAgro, University of Montpellier, France

[9] Laboratory of Forest Biology, Wrocław University, Poland

### Genetic paternity

The file snp_bt.rds[1] includes 908 genotypes (samples) which were successfully sequenced in blue tits. After filtering, 885 samples (884 unique individuals) were kept. Genetic paternity in blue tits was assessed on a total of 109 individual adults (including five recruits, i.e., nestlings in the previous years) and 780 nestlings (thus 884 unique individuals). The file snp_gt.rds[1] includes 1,286 genotypes (samples) which were successfully sequenced in great tits. After filtering, 1,262 samples (1,256 unique individuals) were kept. Genetic paternity in great tits was assessed on a total of 307 individual adults and 949 nestlings.

In blue tits, sampling included 44 broods with social father, 53 broods without social father and 23 broods where no offspring but both parents were sampled (see also below). In great tits, 103 broods with social father were sampled, along with 19 broods without social father and 106 broods where no offspring but both parents were sampled. Broods with no offspring and only parents sampled did not count towards extra-pair statistics, but their genotypes increased sampling groups where social relationships and extra-pair fathers can be established. Among nests where the social father was sampled, genetic analyses identified that in 41% of all cases in blue tits (18/44 nests; 6/40 individual males) and 39% of all cases in great tits (40/103 nests; 40/98 individual males) the social father was cuckolded. This was equivalent to a value of 0.5 social relatedness between males and their social offspring, and GRM relatedness below 0.3 with their social offspring. In blue tits and great tits, 15% and 14% of nestlings (respectively 56/362 and 111/821) were identified as extra-pair (i.e., having 0.5 social relatedness and GRM relatedness below 0.3 with their social father). Among nests where the social father was not sampled, we did not assign individual EPO status to each nestling, but we determined whether such nests contained full or half-siblings (the remaining 418 nestlings in blue tit and 128 nestlings in great tit). In these nests, 54% blue tit nests (28/52, in one nest, there was only one nestling genotyped, so nest status remained not assigned) and 37% great tit nests (7/19) contained half-siblings (i.e., nestlings with 0.5 social relatedness and GRM relatedness between 0.15 and 0.35 with each other). Overall, we observed mixed paternity in 48% of blue tit broods (46/96) and in 39% of great tit broods (47/122). One great tit nestling was classified as an instance of brood parasitism, and three nestlings in a single blue tit nest were not the eggs of the social female (probably a nest take-over). To avoid biases in subsequent analyses on EPP occurrence and proportion of EPO per brood, we filtered out nests where less than 50% of nestlings[2] were successfully genotyped (brood_size), i.e., where brood_size over the initial_brood_size < 0.5 proportion (8 nests in blue tits and 6 in great tits were removed).

### Microsatellite parentage analysis and comparison with SNP data

The DNA of blue tits was extracted using a combination of QIAamp DNA Investigator Kit for adults and QuickGene DNA Tissue Kit for nestlings (Qiagen, Hilden, Germany) and was genotyped at four microsatellite loci: PCA3, PCA7, PCA8, and PCA9[3]. Microsatellite amplification was conducted in separate reactions for each locus. One μL of extracted DNA was used per reaction, and 3 μL for adult samples, which occasionally yielded lower DNA quantities (less blood cells are present in adult superior umbilicus of the shaft). DNA extracted from feathers was amplified in a 25-μL reaction mixture containing 10 pmol of each primer, 10–50 ng of template DNA, 12.5 μL of REDTaq PCR ReadyMix (Sigma-Aldrich), and 7.5 μL of PCR-grade water (Sigma-Aldrich). Forward primers were fluorescently labelled with dyes (Dye2, Dye3, or Dye4; ProOligo). To prevent contamination, all reagents, tubes, and pipettes were exposed to ultraviolet light for 15 minutes before use. Amplification was performed using a Techne Touchgene thermocycler with the following thermal profile: initial denaturation at 94°C for 3 minutes, followed by 34 cycles of 94°C for 1 minute, 55°C for 45 seconds, and 72°C for 45 seconds, and a final cycle at 94°C for 1 minute, 55°C for 45 seconds, and 72°C for 5 minutes. Fragment lengths of the amplified microsatellites were analyzed using a CEQ8000 Beckman Coulter automated sequencer (Beckman Coulter, Fullerton, CA, USA). Data was interpreted using Beckman Coulter Fragment Analysis Software (v. 9.0).

With microsatellites, 14.6% (124/848) of nestlings were identified as extra-pair, and 48% (48/99) of broods were of mixed-paternity (see raw data file "microsats.csv"[1]). 42/90 individual males from 47 nests were identified as cuckolded fathers, meaning their social nest contained at least one EPO. 20 males were identified as extra-pair fathers in 23 nests. Out of 48 nests identified as containing EPO by microsatellites, 43 were confirmed by SNP analysis (two were not sequenced and three were inconsistent regarding extra-pair paternity occurrence). SNP analysis was able to pinpoint the detailed reasons for inconsistencies in all these nests: a) In the first nest, there was no extra-pair paternity but three chicks were eggs laid by another female and not the social mother; b) in the second nest, only five chicks were successfully sequenced - full siblings, but it is possible that the remaining three were the extra-pair ones; c) in the third nest, the social father was not sequenced and all eight nestlings identified by microsatellites as extra-pair were full siblings. Among nests where the social father was not sampled, if all chicks were full siblings, the nests were classified as single paternity nests by the SNP analysis, whereas in the microsatellite analysis if any of the markers did not accord with other social siblings within the nest the nestling was classified as extra-pair and nest as mixed-paternity nest. Furthermore, there were three nests that were identified as mixed-paternity broods by SNPs, while the microsatellite analysis did not. Overall, the occurrence of mixed paternity in a brood was highly consistent between the methods (McNemar's test comparing EPP detected by microsatellites vs. SNPs: $\chi^2 = 0$, df = 1, p = 1), and the relationship between the number of offspring assigned as extra-pair per brood was strong between the methods (Spearman's rho = 0.816, $p < 0.0001$).

### Phenological and environmental variables

*The variables are classified in sets of highly correlated variables (Figs. S2 and S3) describing similar phenological and environmental attributes. Due to the limited number of observations, to prevent multicollinearity, reduce model overfitting and improve the reliability of model selection, we retained as a predictor only one variable within each set that was recorded in the maximum number of nests (here underscored within each set). Fledging age was used as a response variable in separate models (Tables S4 and S5). It was not possible to record all variables in each nest.*

***SET#1:*** **<u>Lay date</u>**, laying date of the first egg, recorded as the number of days from April 1st. First egg dates were estimated by assuming one egg was laid daily or by subtracting 25 days for blue tits and 23 days for great tits from hatching dates for nests with unknown clutch sizes. This estimate included an average of 11 days for egg-laying in blue tits and 9 in great tits, and 14 for incubation; **Hatch date**, date of hatching of the first egg, number of days from April 1st. Hatching dates were determined with ±1 day accuracy from direct observations of parental behavior or nest contents. In 8% of broods, dates were back-calculated by subtracting 19 days from fledging dates[4,5]; **Fledge date**, a day in which all chicks left the nest (number of days from April 1st, fledging dates were calculated by revisiting nests with young and visually aging nestlings, while for inaccessible nests, fledging dates were estimated by adding 18 days to observed hatching dates, identified through parental behaviors such as carrying food or removing eggshells. ***SET#2:*** **Clutch size**, egg counts conducted after incubation; **<u>Initial brood size,</u>** generally denotes the number of hatched nestlings, however as in some nests this data was missing, to avoid missing values for modelling, we used available information at the earliest possible stage: clutch size (16/97 and 12/122 cases in blue tits and great tits respectively) and number of fledglings if clutch size was not available (10/97 and 8/122 cases in blue tits and great tits respectively). In two cases for blue tits and two cases for great tits none of the values was available so we assumed initial brood size as 10 for blue tits and 9 for great tits, which are the average number of hatchlings for the population; **Number of hatched** birds, nestlings that successfully hatched within a clutch; **Number of fledged** birds, the number of nestlings that successfully left the nest.

Additionally:

**Fledging age of nestlings** – defines the time nestlings spent in the nest from hatching till fledging (in days, fledging date – hatching date). Around fledging, nests were observed daily from a distance of 15–50 meters, depending on nest location and parental behavior. A nest was deemed successful if no feeding was observed after the young were 18 days old and when parents were later seen provisioning fledglings. Premature cessation of parental activity (no signs of parents for 90 minutes) indicated failure.

*Nest cavity attributes: all measurements were done according to the methodology described in Wesołowski & Rowiński (2012)*[4] *and Maziarz et al. (2016)*[5]*. It was not possible to measure all attributes in each nest.*

***SET#3:*** **Height**, measured at the bottom of entrance above ground level, in all cavities including inaccessible ones, cavity entrance height was visually estimated for cavities up to 10 m, otherwise measured to nearest 1 m using a clinometer[4,5]; **Tree circumference**, measured by measuring tape at breast height; **Tree circumference at nest** measured by measuring tape at the nest level. ***SET#4:*** **Entrance width,** largest horizontal dimension; **Entrance height**, vertical dimension; **Entrance area**, approximated to an ellipse area = π × 1/2 entrance width × 1/2 entrance height. ***SET#5:*** **Depth**, distance from entrance to the nest measured vertically; **Safety distance**, shortest distance from the lower edge of the entrance to the nest cup (often measured diagonally). ***SET#6:*** **Nest width,** size of nest cup measured perpendicularly to nest length; **Nest length,** size of nest cup measured from entrance to the opposite wall; **Nest bottom area,** nest size approximated to an ellipse area = π × 1/2 nest width × 1/2 nest length; **Cavity volume,** nest size approximated to a cylinder volume = bottom area × depth.

Additionally, categorical variable:

**Exposure,** entrance facing one of the eight cardinal and intercardinal directions, categorized into "coolest" (N, NE and NW), "intermediate" (W and E) and "warmest" (S, SE, SW), which best characterizes thermal conditions at these exposures in spring months in Białowieża.

***References***:

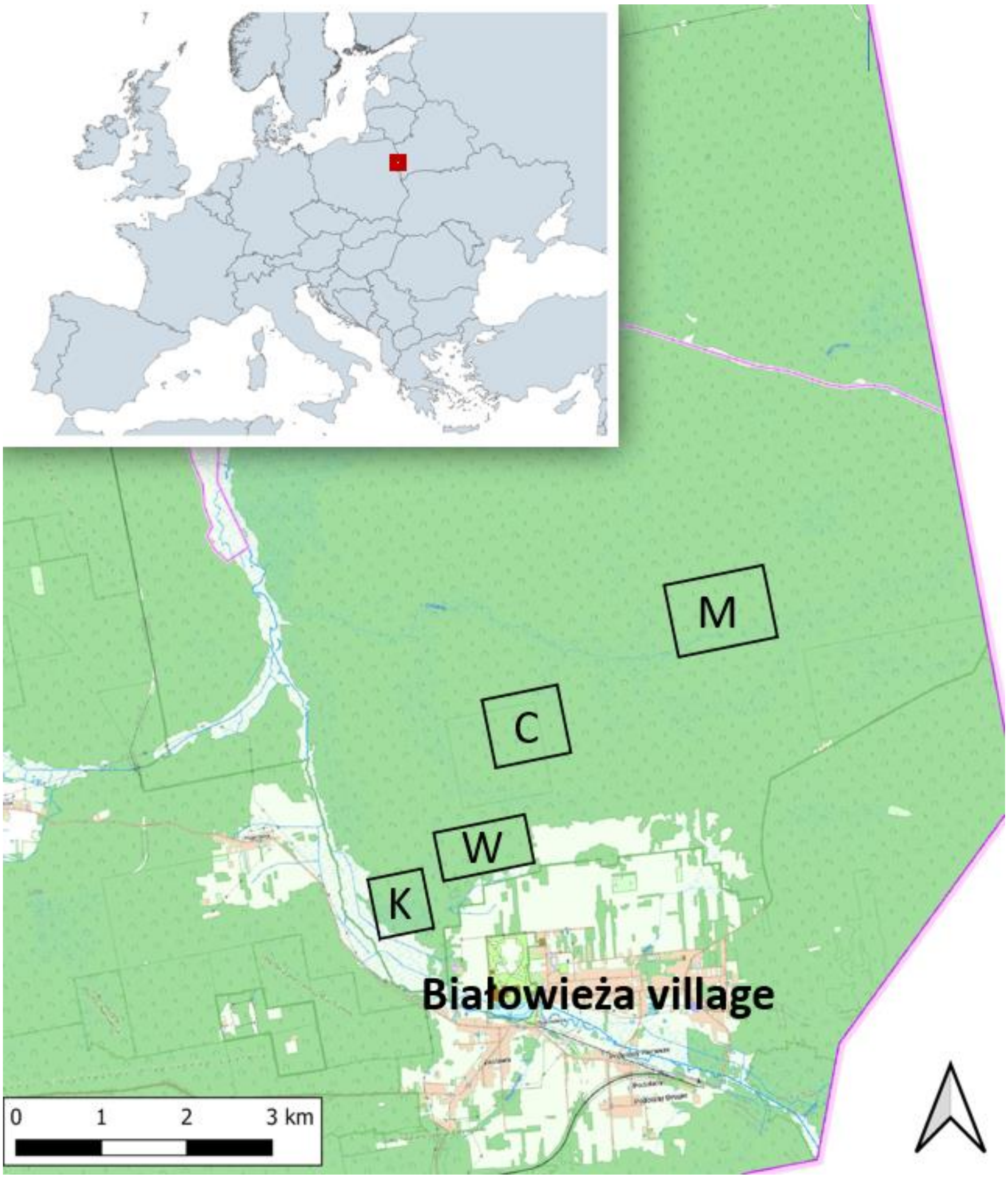

*Figure S1 - Map of study sites within Białowieża National Park*

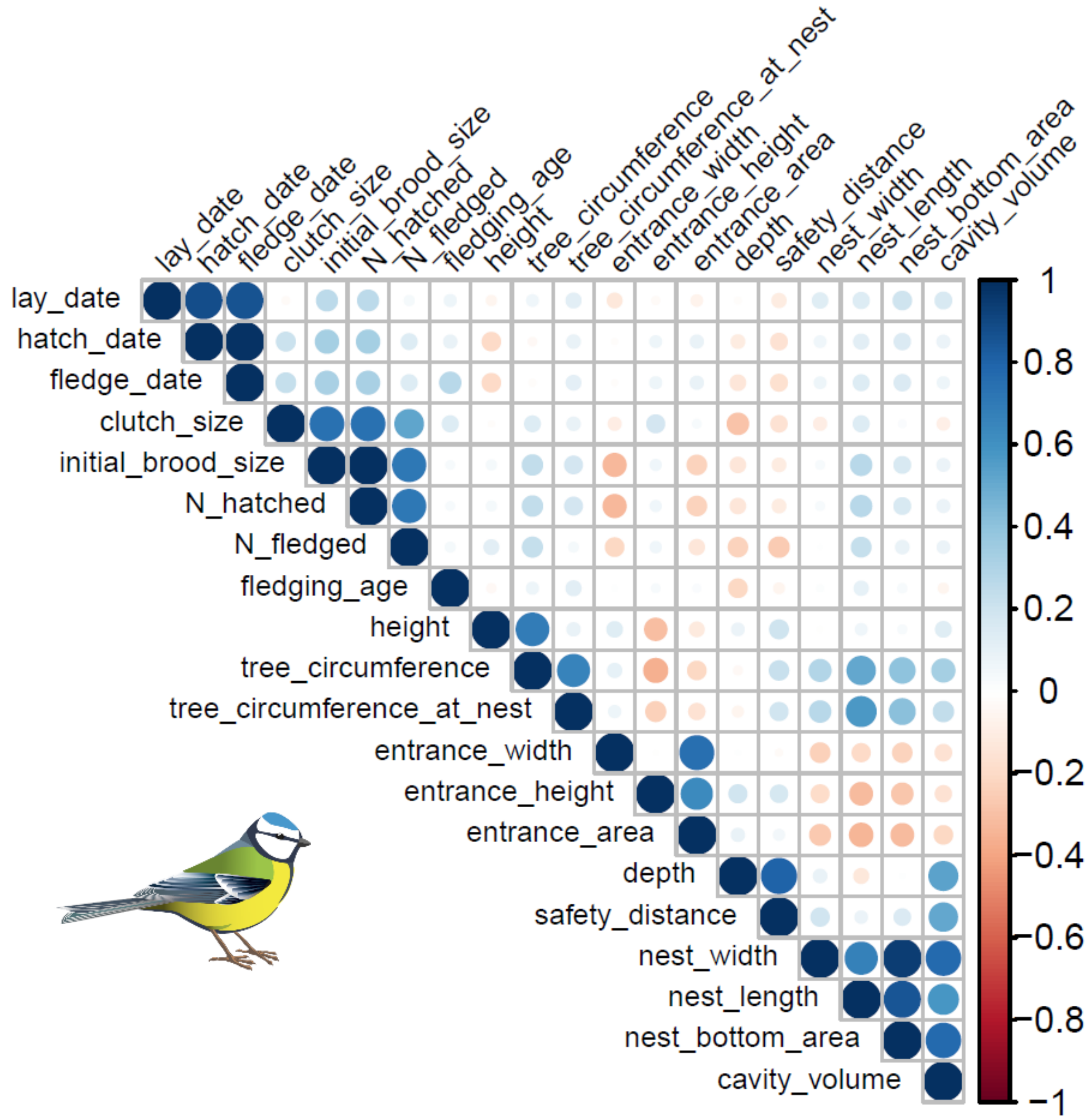

***Figure S2 - Correlation heatmap of the full set of phenological and environmental variables gathered in Białowieża primeval forest blue tit nests***. Blue denotes positive correlations, and red negative. The dot size and color intensity denote correlation strength, with darker colors and larger dots indicating stronger correlations.

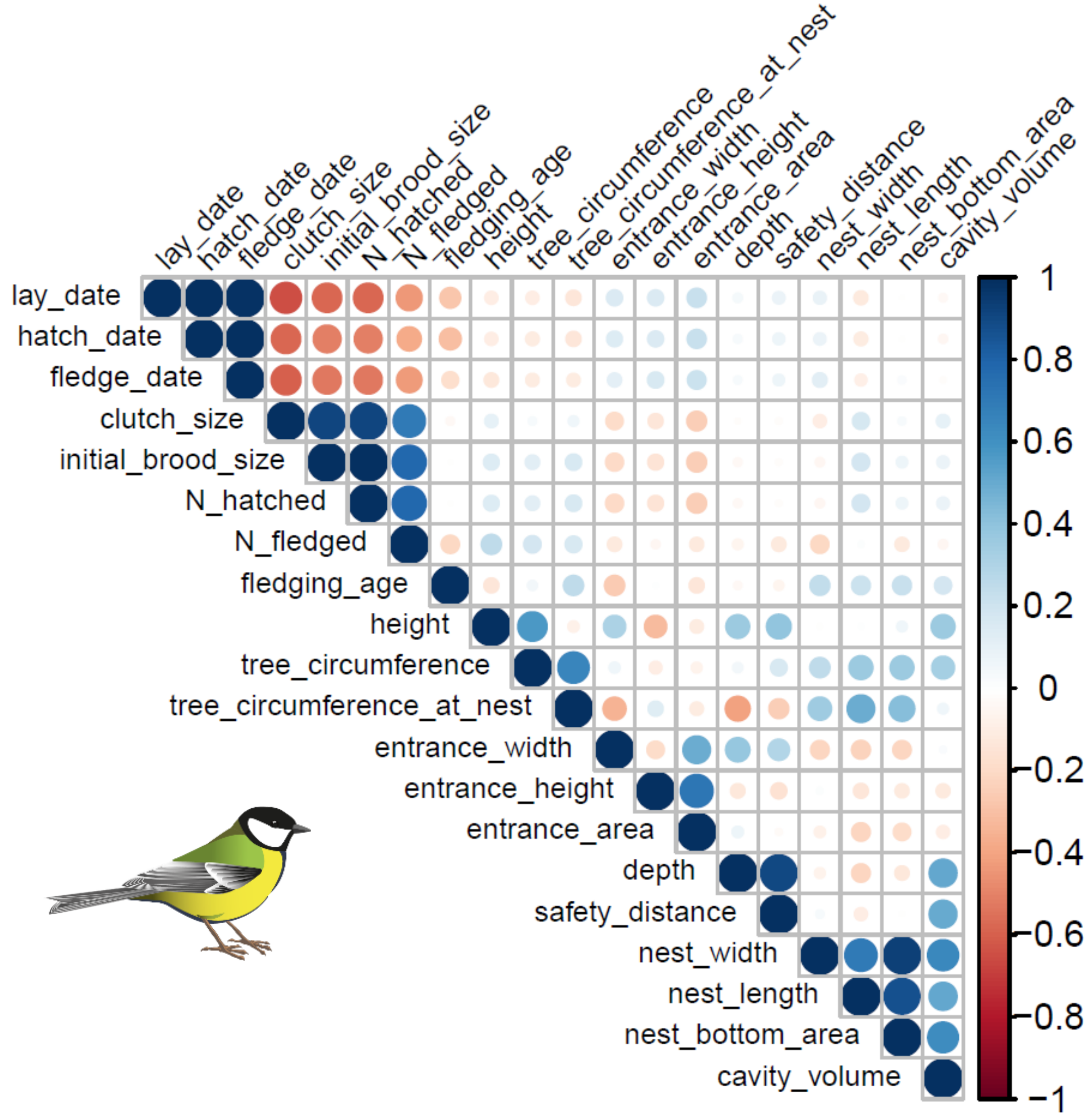

***Figure S3 - Correlation heatmap of the full set of phenological and environmental variables gathered in Białowieża primeval forest great tit nests***. Blue denotes positive correlations and red negative. The dot size and color intensity denote correlation strength, with darker colors and larger dots indicating stronger correlations.

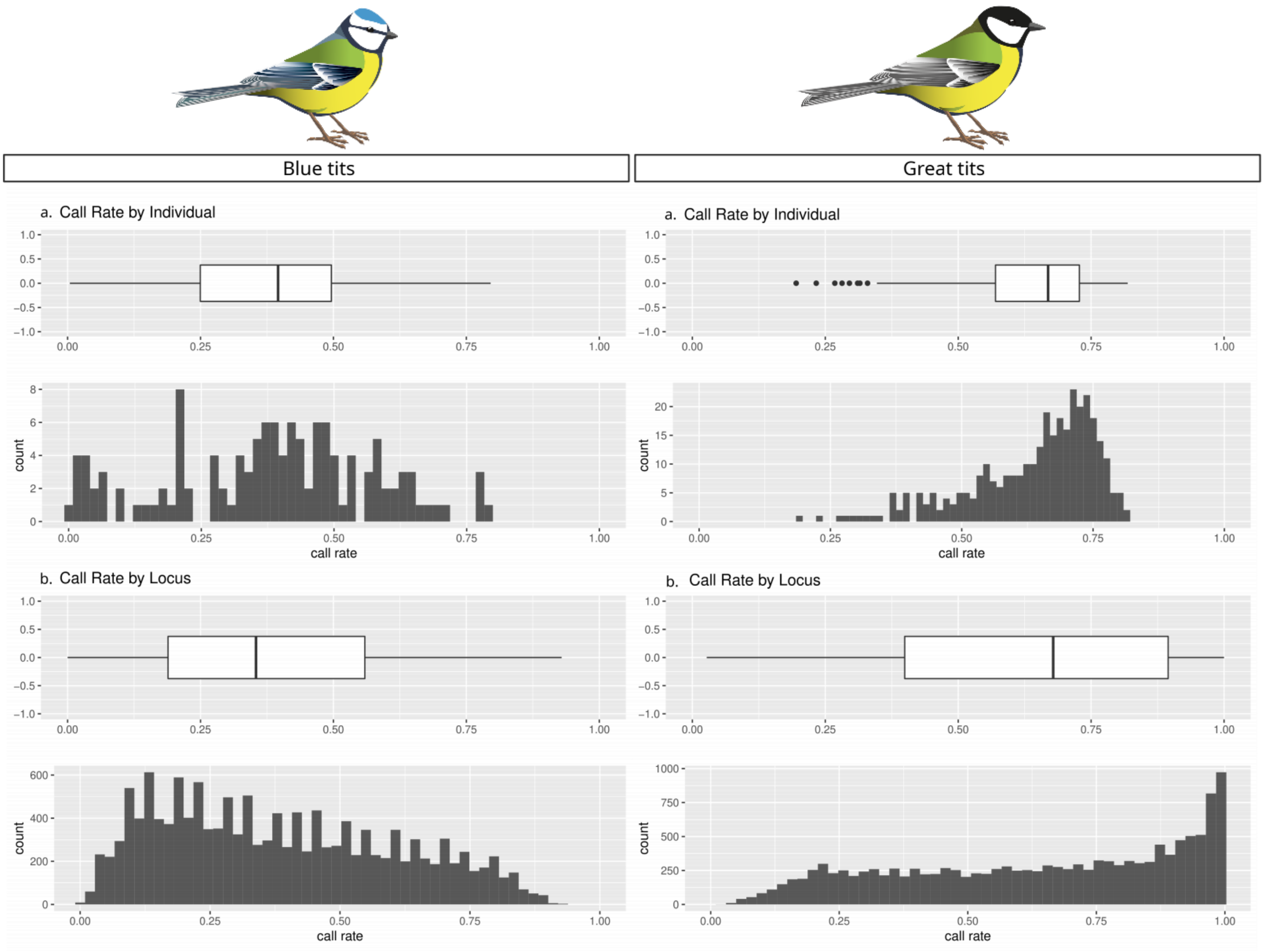

*Figure S4 - Call rate by individual (a) and by locus (b) in adult blue tits and great tits*

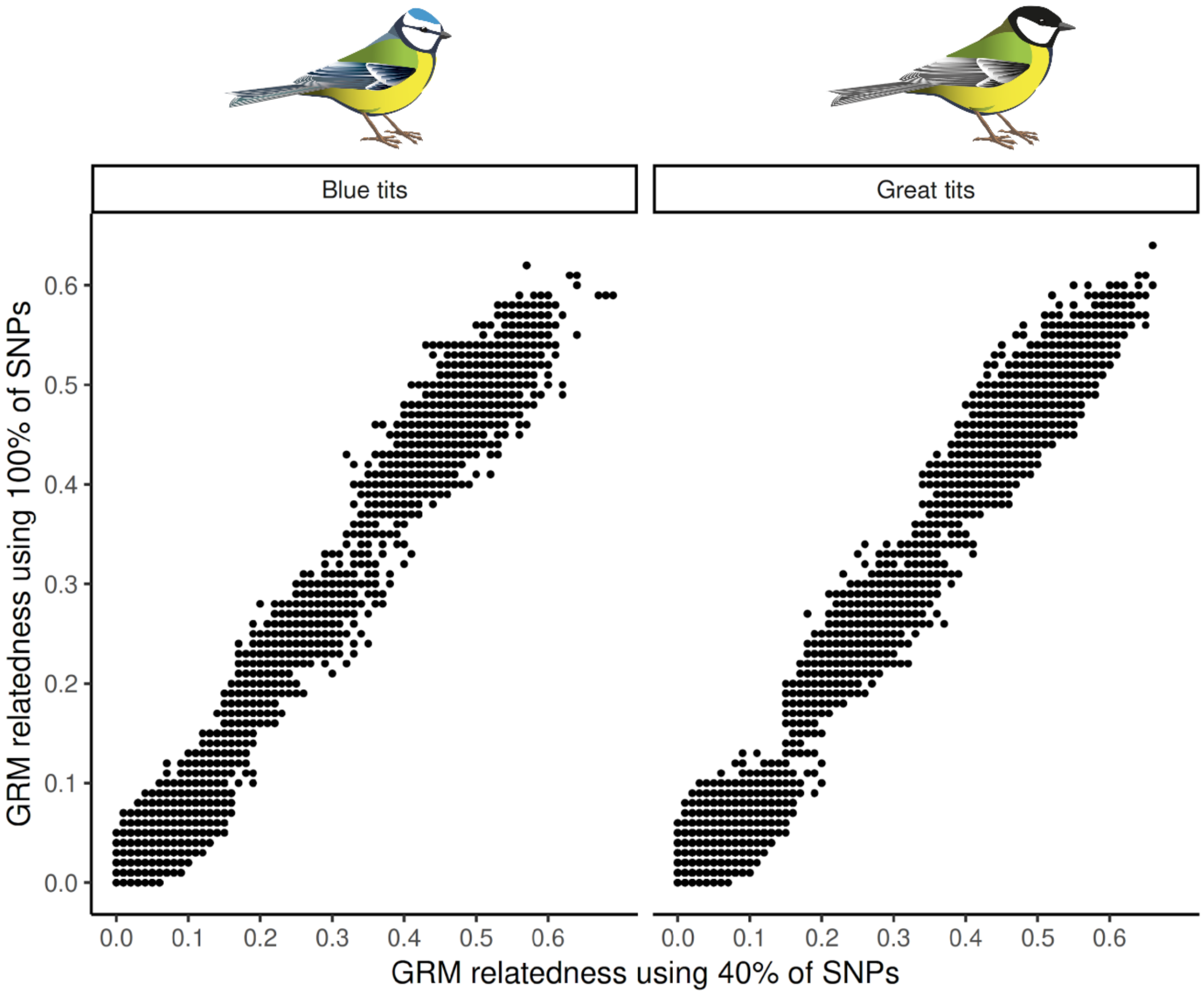

***Figure S5 - Comparison of genetic relatedness calculated with 40% and 100% of the SNP data in blue tits and great tits***

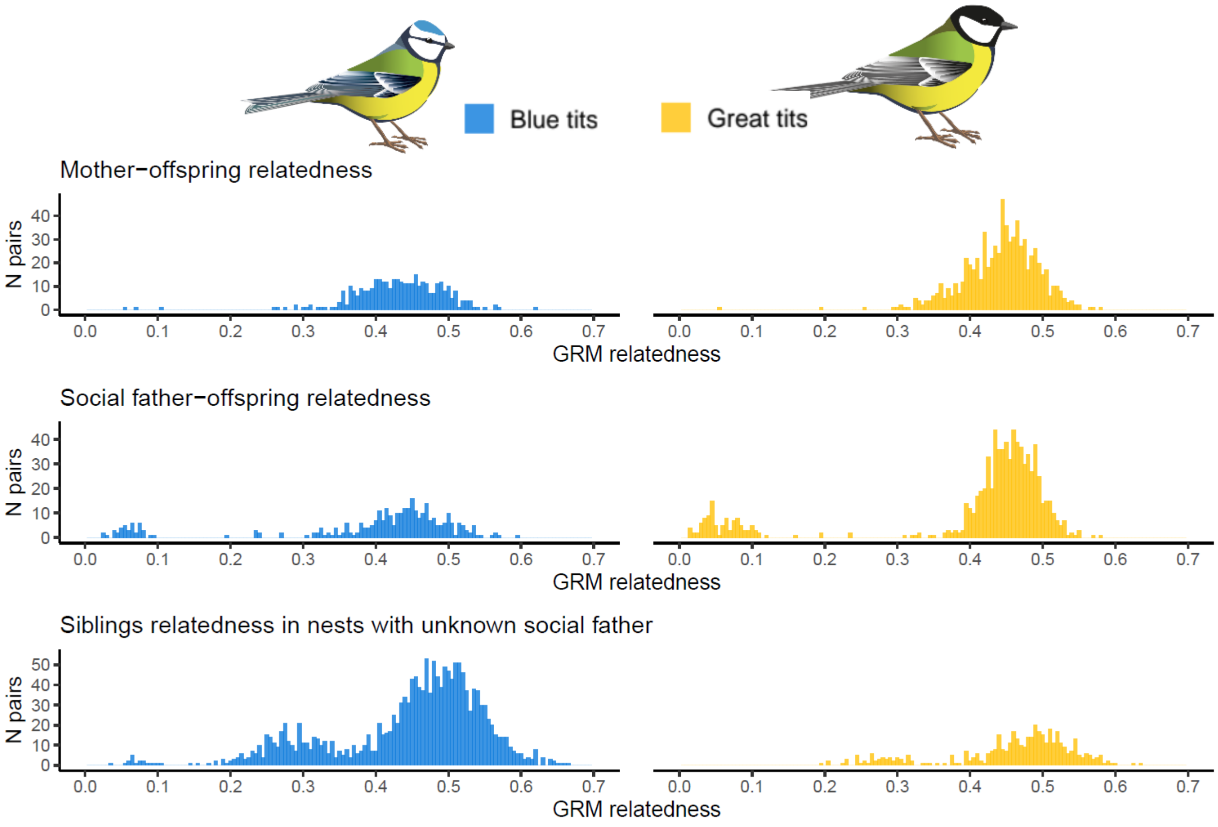

***Figure S6 – a) GRM relatedness values of mothers with their offspring*** for 43 blue tit and 93 great tit nests. The relatedness of mothers to their offspring always reflects the theoretically established coefficient of relatedness r = 0.5, visualized on the GRM with values >0.3. The combination of all relatedness values between mothers and their offspring in the brood results in one peak of relatedness values. In blue tits, there are three cases and in great tits, one case where the mother-offspring relatedness is close to 0; since these four offspring show relatedness values close to 0 with all their respective siblings and their social father, they are with all probability the result of brood parasitism or nest take-over. ***b) GRM relatedness values of social fathers with their offspring*** for 44 blue tit nests and 103 great tits nests. The coefficient of relatedness of social fathers to their offspring is either r=0.5 (visualized on the GRM with values >0.3) or r=0 (i.e., the social father is not the offspring's genetic father, visualized on the GRM with values <0.3, with most values <0.15). The combination of all relatedness values between social fathers and offspring in their broods results in two distinct peaks, reflecting parent-offspring genetic relatedness or a lack of parental relatedness. The handful of cases where GRM relatedness values of social fathers to their offspring range from 0.15 to 0.3 are explained by cases of incest between breeding parents or relatedness between a social father and an extra-pair father. ***c) GRM relatedness of siblings, in nests where the social father was not sampled*** for 53 blue tit nests and 19 great tits nests. The coefficient of relatedness of siblings is either r=0.5 (full-siblings; visualized on the GRM with values >0.3) or r=0.25 (half-siblings; visualized on the GRM with values <0.3). The combination of all relatedness values between siblings results in the presence of two distinct peaks (although less marked than, for instance, between social fathers and offspring), reflecting a full-sibling or half-sibling genetic relatedness. For nests where the social father was not sampled, we did not identify individual extra-pair offspring, but whether extra-pair paternity had occurred in the brood.

*Table S1 - Sample sizes*

| BLUE TIT | individuals collected | | | | individuals succesfully genotyped | | | |
|---|---|---|---|---|---|---|---|---|
| ***year*** | ***N broods*** | ***N nestlings*** | ***N males**** | ***N females**** | ***N broods*** | ***N nestlings*** | ***N males*** | ***N females*** |
| 2005 | 34 | 303 | 55 | 38 | 34 | 303 | 24 | 23 |
| 2006 | 36 | 327 | 57 | 43 | 36 | 321 | 13 | 13 |
| 2007 | 31 | 249 | 45 | 44 | 27 | 156 | 22 | 21 |
| **TOTAL** | **101** | **879** | **157** | **125** | **97** | **780** | **59** | **57** |
| **GREAT TIT** | **individuals collected** | | | | **individuals succesfully genotyped** | | | |
| ***year*** | ***N broods*** | ***N nestlings*** | ***N males**** | ***N females**** | ***N broods*** | ***N nestlings*** | ***N males*** | ***N females*** |
| 2007 | 1 | 8 | 1 | 0 | 1 | 8 | 1 | 0 |
| 2008 | 45 | 327 | 66 | 50 | 45 | 323 | 57 | 42 |
| 2009 | 29 | 223 | 58 | 40 | 29 | 220 | 52 | 29 |
| 2010 | 25 | 217 | 43 | 27 | 25 | 216 | 38 | 25 |
| 2011 | 22 | 183 | 52 | 29 | 22 | 182 | 50 | 28 |
| **TOTAL** | **122** | **958** | **220** | **146** | **122** | **949** | **198** | **124** |

*Includes adults that bred in the cavities and individuals found on territories from winter till the start of season, early spring

Some individuals were sampled more than once across study years, either as nestling or breeding parent.

***Table S2 – Generalized linear models examining the occurrence of extra-pair offspring in the broods of Białowieża primeval forest blue tits and great tits.*** Response variable modelled as absence (0) or presence (1). Predictor variables were selected based on model averaging across all models with ΔAICc <1.5. Exposure "warmest" was the reference category for the fixed predictor. Significant differences (P < 0.05) are in bold. Marginal (R2m) and conditional (R2c) R-squared are shown.

**Occurrence of extra-pair offspring in the brood,** GLM binomial

| | 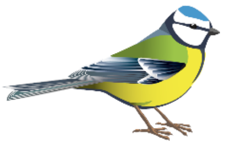 | | | **(n = 80 nests)** | 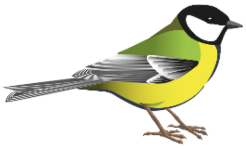 | | | **(n = 99 nests)** |
|---|---|---|---|---|---|---|---|---|
| | **Estimate** | **SE** | **z** | **p** | **Estimate** | **SE** | **z** | **p** |
| **(Intercept)** | -0.94 | 1.67 | -0.56 | 0.576 | 0.41 | 0.80 | 0.51 | 0.611 |
| **depth** | 0.09 | 0.04 | 2.09 | **0.037** | -0.02 | 0.02 | -0.99 | 0.323 |
| **height** | 0.12 | 0.05 | 2.56 | **0.010** | -0.05 | 0.07 | -0.66 | 0.507 |
| **entrance area** | -0.03 | 0.03 | -0.91 | 0.361 | -0.01 | 0.01 | -0.82 | 0.413 |
| **initial brood size** | -0.12 | 0.13 | -0.95 | 0.344 | | | | |
| **exposure (intermediate)** | | | | | 1.55 | 0.59 | 2.63 | **0.009** |
| **exposure (coolest)** | | | | | 0.92 | 0.52 | -1.78 | 0.075 |
| **nest bottom area** | | | | | -0.003 | 0.002 | -1.28 | 0.199 |
| ***Random effects*** | ***Variance*** | | | | ***Variance*** | | | |
| ***year*** | *0.000 (removed)* | | | | *0.000 (removed)* | | | |
| ***plot*** | *0.000 (removed)* | | | | *0.000 (removed)* | | | |
| **$R^2$m / $R^2$c** | 0.253 / 0.253 | | | | 0.143/ 0.143 | | | |

***Table S3 - Generalized linear (mixed) models examining the proportion of extra-pair offspring per brood in the broods of Białowieża primeval forest blue tits and great tits.*** Predictor variables were selected based on model averaging across all models with ΔAICc <1.5. Exposure "warmest" was the reference category for the fixed predictor. Significant differences (P < 0.05) are indicated in bold. Marginal (R2m) and conditional (R2c) R-squared are shown.

**Proportion of extra-pair offspring per brood,** GLMM binomial

| | 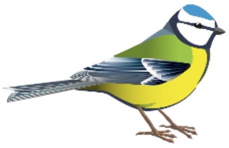 (n = 37 nests) | | | | 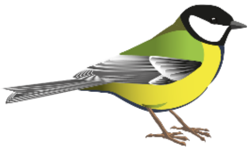 (n = 83 nests) | | | |
|---|---|---|---|---|---|---|---|---|
| | **Estimate** | **SE** | **z** | **p** | **Estimate** | **SE** | **z** | **p** |
| **(Intercept)** | -5.52 | 1.13 | -4.88 | **<0.0001** | 2.11 | 0.79 | 2.68 | **0.007** |
| **depth** | 0.08 | 0.02 | 3.18 | **0.001** | -0.03 | 0.01 | -2.61 | **0.009** |
| **height** | 0.09 | 0.03 | 3.13 | **0.002** | -0.02 | 0.03 | -0.66 | 0.509 |
| **lay date** | 0.07 | 0.04 | 1.74 | 0.082 | | | | |
| **entrance area** | | | | | -0.02 | 0.01 | -2.41 | **0.016** |
| **initial brood size** | | | | | -0.26 | 0.07 | -3.76 | **0.0002** |
| **exposure (intermediate)** | | | | | 0.46 | 0.27 | 1.67 | 0.095 |
| **exposure (coolest)** | | | | | -0.10 | 0.27 | -0.36 | 0.714 |
| **nest bottom area** | | | | | -0.001 | 0.001 | -1.11 | 0.267 |
| ***Random effects*** | *Variance* | | | | *Variance* | | | |
| ***year*** | *0.114* | | | | *0.000 (removed)* | | | |
| ***plot*** | *0.000 (removed)* | | | | *0.000 (removed)* | | | |
| **$R^2$m / $R^2$c** | 0.165 / 0.193 | | | | 0.136 / 0.136 | | | |

***Table S4 – Generalized and general linear (mixed) models examining variation in a) number of fledged young and b) fledging age*** (i.e. time spent by the nestlings in the nest from hatching till fledging and a measure of parental investment) in the broods of Białowieża primeval forest blue tits and great tits. Predictor variables were selected based on model averaging across all models with ΔAICc <1.5 for number of fledged young and fledging age in great tits and the best models are presented for both responses in blue tits (since just one model's ΔAICc <1.5), where epp occurrence was always retained as a fixed factor of interest. Epp occurrence "0" (no extra-pair offspring in a nest) and exposure "warmest" were the reference categories for the predictors. Significant differences ($P < 0.05$) are indicated in bold. Marginal (R2m) and conditional (R2c) R-squared are shown.

### A) Number of fledged young, LMM for blue tits and GLM poisson for great tits

| | 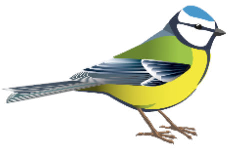 | | | **(n = 78 nests)** | 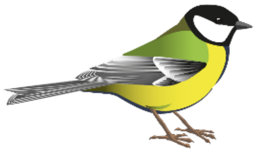 | | | **(n = 98 nests)** |
|---|---|---|---|---|---|---|---|---|
| | **Estimate** | **SE** | **t** | **p** | **Estimate** | **SE** | **z** | **p** |
| **(Intercept)** | -0.13 | 1.43 | -0.09 | 0.928 | 0.70 | 0.31 | 2.30 | **0.021** |
| **epp occurrence (1)** | 0.02 | 0.54 | 0.05 | 0.964 | 0.14 | 0.08 | 1.79 | 0.074 |
| **initial brood size** | 0.88 | 0.13 | 6.57 | **<0.0001** | 0.14 | 0.03 | 5.46 | **<0.0001** |
| **lay date** | | | | | -0.006 | 0.003 | -1.51 | 0.131 |
| **height** | | | | | 0.01 | 0.01 | 1.18 | 0.238 |
| ***Random effects*** | *Variance* | | | | *Variance* | | | |
| ***year*** | *0 (removed)* | | | | *0 (removed)* | | | |
| ***plot*** | *0.099* | | | | *0 (removed)* | | | |
| **$R^2m$ / $R^2c$** | 0.361/ 0.361 | | | | 0.432 / 0.432 | | | |

### B) Fledging age, LMM

| | 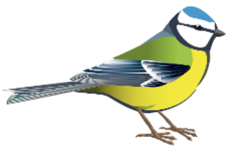 | | | **(n = 76 nests)** | 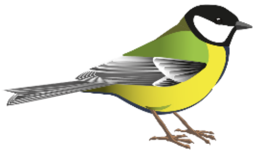 | | | **(n = 80 nests)** |
|---|---|---|---|---|---|---|---|---|
| | **Estimate** | **SE** | **t** | **p** | **Estimate** | **SE** | **t** | **p** |
| **(Intercept)** | 19.49 | 0.15 | 131.80 | **<0.0001** | 17.70 | 0.41 | 43.00 | **<0.0001** |
| **epp occurrence (1)** | -0.04 | 0.20 | -0.19 | 0.853 | 0.09 | 0.37 | 0.23 | 0.816 |
| **exposure (intermediate)** | | | | | 0.69 | 0.47 | 1.48 | 0.143 |
| **exposure (coolest)** | | | | | 0.78 | 0.42 | 1.85 | 0.068 |
| ***Random effects*** | *Variance* | | | | *Variance* | | | |
| ***year*** | *0 (removed)* | | | | *0 (removed)* | | | |
| ***plot*** | *0.012* | | | | *0.209* | | | |
| **$R^2m$ / $R^2c$** | 0.0005 / 0.016 | | | | 0.048/ 0.122 | | | |

***Table S5 - Generalized and general linear (mixed) models examining variation in a) number of fledged young and b) fledging age*** (i.e. time spent by the nestlings in the nest from hatching till fledging) in the broods of Białowieża primeval forest blue tits and great tits. Predictor variables were selected based on model averaging across all models with ΔAICc <1.5 for number of fledged young in blue tits and great tits and the best models are presented for fledging age in blue tits and great tits (since just one model's ΔAICc <1.5), where proportion of EPO per nest was always retained as fixed factor of interest. Significant differences ($P < 0.05$) are indicated in bold. Marginal ($R^2m$) and conditional ($R^2c$) R-squared are shown.

### A) Number of fledged young, LM for blue tits and GLM poisson for great tits

| | 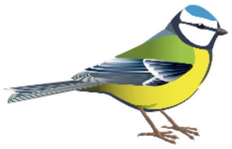 | | | **(n = 36 nests)** | 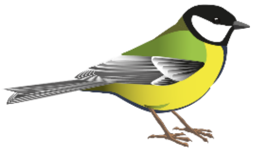 | | | **(n = 83 nests)** |
|---|---|---|---|---|---|---|---|---|
| | **Estimate** | **SE** | **t** | **p** | **Estimate** | **SE** | **z** | **p** |
| **(Intercept)** | 10.90 | 3.14 | 3.47 | **0.002** | 0.51 | 0.27 | 1.89 | 0.058 |
| **proportion epo** | -3.19 | 1.74 | -1.84 | 0.076 | -0.05 | 0.16 | -0.34 | 0.737 |
| **initial brood size** | 0.52 | 0.19 | 2.70 | **0.011** | 0.16 | 0.03 | 6.27 | **<0.0001** |
| **lay date** | -0.18 | 0.09 | -2.15 | **0.040** | | | | |
| **height** | | | | | 0.01 | 0.01 | 1.17 | 0.242 |
| **nest bottom area** | -0.01 | 0.007 | -1.87 | 0.071 | -0.0006 | 0.0003 | -1.67 | 0.094 |
| **entrance area** | -0.08 | 0.04 | -2.21 | **0.035** | | | | |
| ***Random effects*** | *Variance* | | | | *Variance* | | | |
| ***year*** | *0 (removed)* | | | | *0 (removed)* | | | |
| ***plot*** | *0 (removed)* | | | | *0 (removed)* | | | |
| **$R^2m$ / $R^2c$** | 0.443 / 0.443 | | | | 0.388/ 0.388 | | | |

### B) Fledging age, LMM

| | 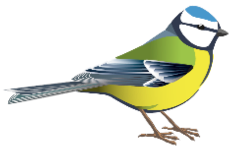 | | | **(n = 35 nests)** | 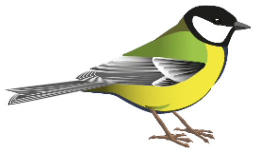 | | | **(n = 68 nests)** |
|---|---|---|---|---|---|---|---|---|
| | **Estimate** | **SE** | **t** | **p** | **Estimate** | **SE** | **t** | **p** |
| **(Intercept)** | 19.36 | 0.23 | 82.58 | **0.0001** | 18.00 | 0.33 | 54.51 | **<0.0001** |
| **proportion epo** | -0.07 | 0.64 | -0.11 | 0.915 | 0.61 | 0.70 | 0.87 | 0.386 |
| ***Random effects*** | *Variance* | | | | *Variance* | | | |
| ***year*** | *0.078* | | | | *0 (removed)* | | | |
| ***plot*** | *0 (removed)* | | | | *0.181* | | | |
| **$R^2m$ / $R^2c$** | 0.0003 / 0.111 | | | | 0.011 / 0.081 | | | |
